# Transient interactions and influence among bacteria in field-grown *Arabidopsis thaliana* tissues

**DOI:** 10.1101/2020.03.05.974105

**Authors:** Kathleen Beilsmith, Matthew Perisin, Joy Bergelson

## Abstract

Interactions between bacteria are thought to play an important role in the assembly of plant microbial communities (1), yet the extent of temporal variation in these interactions is unclear. We inferred interactions from sequence-based counts of bacteria in a series of *Arabidopsis thaliana* tissue samples spanning major developmental transitions in the plant life cycle (2). Bacterial interactions were transient, even among variants found together at consecutive developmental stages. The overwhelming majority of these interactions were positive, indicating that competition for the plant niche might be a less important driver of bacterial abundances than cooperation or common responses to host and environmental factors. Over time, interaction networks diverged from an initial scale-free structure and became increasingly modular. In all networks, we found evidence of a hierarchical structure in which hub bacteria bridged network modules. However, the identities of bacteria in these influential roles also varied during plant growth.

## Main

Microbial communities undergo shifts in diversity and composition as seasons change and their hosts mature (3,4). However, temporal variation of microbe-microbe interactions within these communities has not been characterized. Since most natural surveys of plant microbial communities take snapshots of a single life stage, it is difficult to assess whether the inferred interactions, especially those of highly influential “hub” bacteria, consistently shape the community or are subject to turnover as the host ages.

To characterize this temporal variation, we inferred bacterial interactions in tissue samples spanning the vegetative, flowering, and senescent phases of an annual plant’s development. Over two years, the bacteria inhabiting *A. thaliana* were surveyed by amplifying and sequencing a portion of the 16S rRNA gene (16S) in root and phyllosphere samples from plants grown at two sites in southeast Michigan: Michigan State Extension Center (ME) and Warren Woods Ecological Field Station (WW). Amplicon sequence variants (ASVs) defined the bacterial lineages present in each sample, albeit with limited resolution below the family level (5). Using the SPIEC-EASI pipeline (6), interactions were inferred between ASVs based on an inverse covariance matrix generated from transformed counts. We assigned direction and magnitudes to the interactions based on abundance correlations (7) between the ASVs (<u>Supplementary Methods</u>).

A previous study of *A. thaliana* leaf microbial networks reported that most (86.5%) interactions between bacteria were positive (8). We too found that the abundance correlations were overwhelmingly positive throughout development, regardless of the tissue sampled or the site of harvest (Table 1).

**Table 1.**
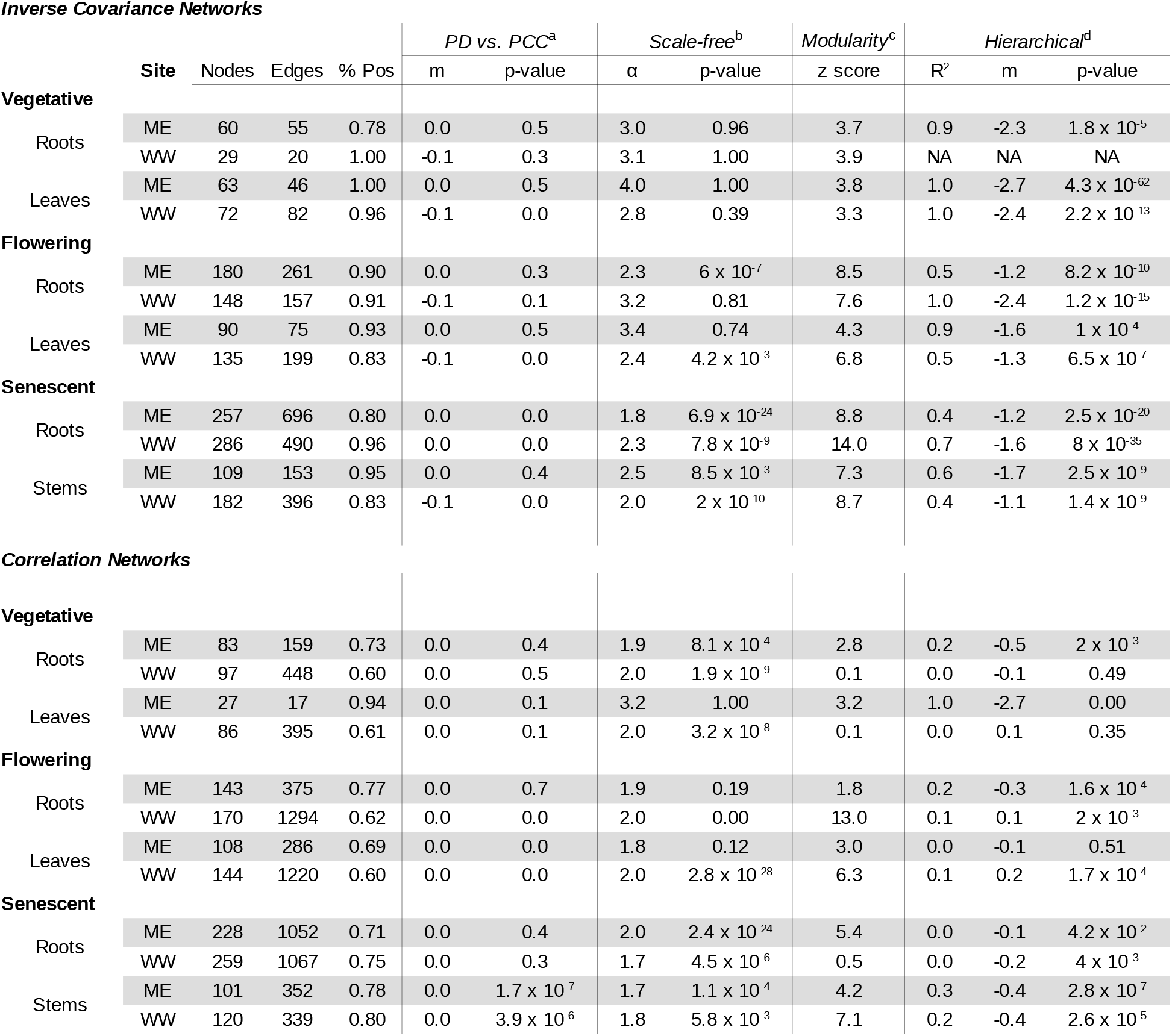
Properties of bacterial interaction networks from *Arabidopsis thaliana*. The top panel presents results from networks inferred with an inverse covariance approach and the bottom panel presents results from correlation-based networks. Rows are divided by stage, tissue, and field site. The first three columns present the number of bacteria (nodes), interactions (edges), and the fraction of positive interactions (% Pos) in each network. Further right, columns present results for four patterns in network structure: (a) The slope (m) and F-test p-value for a linear model fit to phylogenetic distance (PD) vs. Pearson correlation coefficient (PCC); (b) α and Kolmogorov-Smirnov p-value for a power law fit to the network degree distribution Pr(k) ∝ k^−α^; (c) a z-score for the empirical network modularity in a distribution obtained by simulating random networks of the same size; and (d) the coefficient of determination (R^2^), slope (m) and F-test p-value for a linear model fit to log10 local clustering coefficient C(k) vs. log10 degree k.

Interactions are visualized for root networks in Figure 1A and for phyllosphere networks in <u>Figure S1</u>.

**Figure 1.**
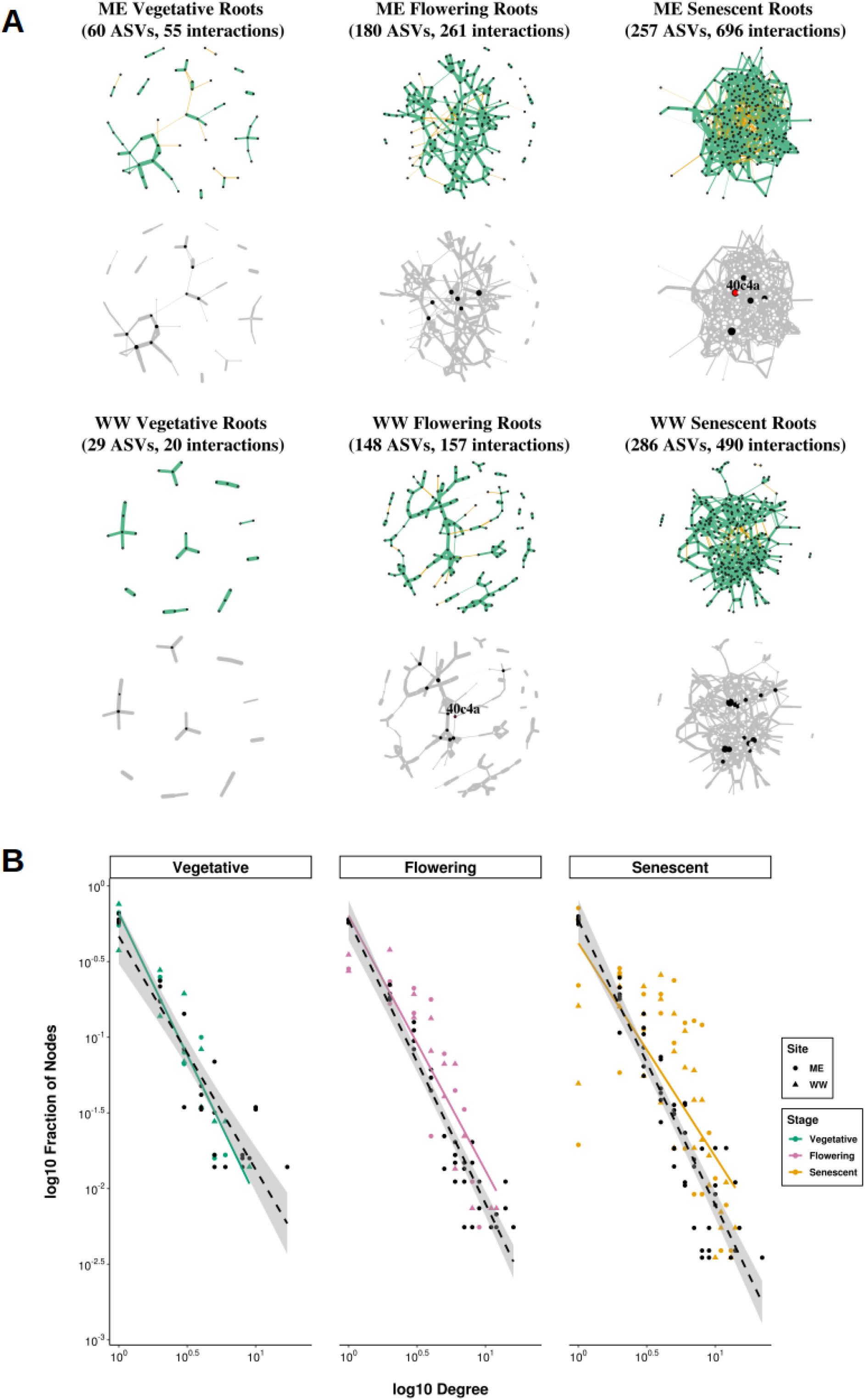
Bacterial interactions in *A. thaliana* remain overwhelmingly positive while hub identities and network structure change during development. (A) The inferred interaction networks for root bacteria at each stage (column) and site (row) are displayed. In the top image, color indicates positive (green) and negative (orange) abundance correlations between 16S ASVs in the network. The thickness of the edges between ASVs represents the absolute value of the correlation coefficient. While networks inference from sequencing data cannot distinguish between direct microbe-microbe interactions and those mediated by, for example, the plant host, patterns can be observed in the nature of the interactions and the ASV involved in them. At all developmental stages, a majority of interactions are positive. The bottom image highlights the network’s hub bacteria, as determined by degree and betweenness centrality above the ninetieth percentile, in black. Hub identities are inconsistent: only one bacterial variant (40c4a, colored red) in the genus *Geodermatophilus* is classified as a hub in multiple root networks. (B) The relationship between degree and the fraction of nodes with that degree is plotted on a log10 scale for all nodes in plant root and phyllosphere networks. The empirical data are distinguished by field site (shape) and developmental stage (color) and plotted against a null distribution (black) generated for networks of the same size with a Barabási-Albert model (19). As shown by best-fit lines with a 95% confidence interval for the null distribution, the relationships increasingly depart from the expectation for a scale-free network as development progresses.

The high fraction of positive relationships suggests that competitive interactions do not stabilize plant bacterial communities as they do in models of the human gut microbiome (9). A preponderance of positive interactions is perhaps not surprising given that we considered bacteria only; broader surveys of the plant microbiome indicate that negative interactions are more common between kingdoms than within them (8,10). Positive correlations between bacterial ASV abundances might arise from metabolic cooperation (11), a confounding non-bacterial microbe (10) or a confounding abiotic factor (12). Irrespective of the cause, we found that the strength of these inferred ecological relationships was unrelated to the phylogenetic distances between bacteria (estimated by Pearson correlation coefficients (PCC) and phylogenetic branch length (PD) respectively) (Table 1), although we note that 16S has poor resolution in distinguishing closely related species.

Bacterial interactions in the roots and the phyllosphere were transient, at least with respect to our relatively coarse temporal sampling. With each successive developmental transition in roots, 60 to 80% of bacteria present before the transition were retained. These bacteria increased their number of interaction partners (<u>Figure S2</u>), but less than 10% of their previous interactions remained intact (<u>Figure S3</u>). In leaves, a smaller fraction of the community was conserved across the transition from vegetative growth to flowering. Of bacteria that entered the network during vegetative growth, 35% to 51% were present after flowering. Again, less than 10% of their relationships were conserved across stages. No more than 30% of interactions were conserved across any two networks within a tissue type, developmental stage, or field site.

The influence of bacteria was also inconsistent across tissues, sites, and developmental stages. Hub bacteria were identified by degree and betweenness centrality scores above the ninetieth percentile for nodes in a network. Neither of these metrics correlated with the prevalence or abundance of the bacteria in the dataset (<u>Figure S4</u>). No bacteria were consistently designated as hubs throughout plant life and only 11% persisted as hubs across any tissues, sites or more than one stage (<u>Figure S5</u>). For example, only one recurring hub was observed in root networks (Figure 1A). The inconsistency of influential bacteria was robust to lowering the thresholds on degree and betweenness centrality that defined hubs (<u>Table S1</u>). Hub microbes are proposed to propagate the effects of the abiotic environment or host plant throughout the microbial community (8,13). Although hub status in computed networks is insufficient to establish keystone taxa (14), the inconsistency we find in hub identities indicates that the microbes playing keystone roles in plant microbial communities could shift over a host’s lifetime.

Networks from all tissues and both sites displayed a power-law relationship between node clustering coefficient and degree throughout development, consistent with a hierarchical structure in which hubs connect modules of lower-degree nodes (15). Other structural characteristics of the networks were in flux as communities assembled during plant growth. We tested whether the degree distribution of each network was well fit by the power-law distribution characteristic of scale-free networks (16, 17). Although early networks in roots and leaves were scale-free, some flowering networks and all senescent networks diverged from this structure (Table 1, Figure 1B), adding to evidence that strict scale-free structure is uncommon in biological networks (18) and showing that systems can move in and out of this paradigm over time. As networks became less scale-free, they became more modular relative to networks of the same size with randomized edges (Table 1). Modularity is linked to the stability of ecological systems. High modularity destabilizes communities, especially when interactions are strongly positive (19).

We inferred these interaction networks with an inverse covariance approach intended to minimize spurious connections in sparse networks. To determine whether our results were robust to network inference method (20), we repeated the analysis with networks inferred using only the correlations of bacterial abundances and thresholds for their p-values and magnitudes (Table 1). In correlation-based networks, interactions remained overwhelmingly positive and unrelated to the phylogenetic distance between bacteria. Interactions remained transient, with an average of only 10 to 25% of interactions shared between networks within a tissue, stage, or site. The identities of influential bacteria were again transient across space and time, with less than 30% of hubs holding that status in more than one network (<u>Figure S6</u>). However, these correlation-based networks lacked the evidence of hierarchy and the directional temporal trends in modularity and structure that were observed in the inverse covariance networks. Structural features of the correlation networks varied as much within developmental stages as between them, perhaps because the addition of spurious edges blurred biological patterns.

## Data

The data used in this study can be accessed in the NCBI’s Sequence Read Archive, BioProject ID PRJNA607544 (available March 31, 2020). The community count table, sample metadata, taxonomy, and 16S tree used in the analysis were generated by Beilsmith et al (2). These files and the scripts needed to reproduce the analysis are available at: https://github.com/krbeilsmith/KBMP2020_Networks

## Acknowledgements

KB and MP were supported by the University of Chicago Biological Sciences Division and by NIH T32 GM 7197. MP was supported by a Department of Education GAANN fellowship. Research was supported by NIH grant R01 GM 083068 and James S. McDonnell Foundation 220020237. The authors thank members of the Bergelson lab and the Department of Ecology and Evolution at the University of Chicago.

## Supplementary information

### Supplementary Methods

Beilsmith2020_SupplementaryMethods.doc

<u>Figure S1</u>

These phyllosphere bacterial networks further support points made with the root networks in Figure 1A. Color indicates positive (green) and negative (orange) abundance correlations and line thickness corresponds to the correlation coefficient. Hubs are highlighted in black and those conserved across networks are red and labeled with partial ASV identifiers. Only one ASV (e8ab0), in the genus *Nocardioides*, was conserved.

Beilsmith2020_SupplementaryFigure_S1.tif

<u>Figure S2</u>

The number of ASV interaction partners tends to increase over major developmental transitions. For ASVs (nodes) remaining after a transition, the number of edges (degree) before and after the transition were compared. On the x-axis, positive numbers indicate net connections gained while negative numbers indicate net connections lost. The bar height on the y-axis indicates the frequency of conserved nodes with the corresponding number of gains or losses. Interactions gained and lost are shown for the transition to flowering in roots and rosette leaves and for senescence in roots (vertical panels).

Beilsmith2020_SupplementaryFigure_S2.tif

<u>Figure S3</u>

Interactions in the networks are transient, as evidenced by the low number of edges conserved between ASVs that occur before and after major developmental transitions. Network comparisons are indicated on the x-axis by filled circles connected by lines. The number of overlapping edges in each comparison is shown on the y-axis. The total number of edges in each network is plotted to the left of the network name.

Beilsmith2020_SupplementaryFigure_S3.tif

<u>Figure S4</u>

Hubs were identified by degree and betweenness centrality. These criteria for influence did not correlate with ASV prevalence (the number of samples in which the ASV was present) or raw abundance (the total number of counts for the ASV).

Beilsmith2020_SupplementaryFigure_S4.tif

<u>Figure S5</u>

Distributions of recurring hubs (columns) in inverse covariance networks. Hubs are organized by family and compared across developmental stages, field sites (vertical panels) and tissues (rows). Black cells indicate the ASV is both present and meets the hub criteria; white cells indicate the ASV is either not present or not a hub.

File type: TIFF (.tif)

File name: Beilsmith2020_SupplementaryFigure_S5.tif

<u>Figure S6</u>

Distributions of recurring hubs (columns) in correlation-based networks. Hubs are organized by family and compared across developmental stages, field sites (vertical panels) and tissues (rows). Black cells indicate the ASV is both present and meets the hub criteria; white cells indicate the ASV is either not present or not a hub.

File type: TIFF (.tif)

File name: Beilsmith2020_SupplementaryFigure_S6.tif

<u>Table S1</u>

The transience of influence in networks is demonstrated by the inconsistency between them in the ASVs designated as putative hubs. Only 11% of hubs and 30% of hubs were conserved between at least two inverse covariance and correlation-based networks, respectively. When the criteria for hubs were lowered to the eightieth percentile of degree and betweenness centrality for nodes in a network, these fractions did not greatly increase. Even when the thresholds were lowered to the seventieth percentile for hub metrics, only 21% and 37% of hubs recurred across at least two inverse covariance and correlation-based networks, respectively.

Beilsmith2020_SupplementaryTable_S1.xls

