## Supplementary Methods for "Transient interactions and influence among bacteria in field-grown *Arabidopsis thaliana* tissues"

Dataset and software

Plant tissue harvest, extraction, and sequence data processing are described in Perisin 2016 (1) and Beilsmith, Perisin, and Bergelson 2020 (2). The 16S rRNA gene (16S) tree, DADA2-generated ASV count table, QIIME-SILVA (release 128) taxonomy, and metadata from that study were used to construct a phyloseq (3) object in R (4) for analysis. The R commands that reproduce the analysis described below are available at <https://github.com/krbeilsmith/KBMP2020_Networks>. Networks were visualized and analyzed, including randomizations and modularity calculations, with igraph (5). Phylogenetic distances between bacterial ASVs were added to the networks with vegan (6). Figures and supplemental figures were produced with igraph, ggplot2 (7), and ggpubr (8).

Sample filtering and classification

Using phyloseq, the dataset was filtered to exclude samples from soil, flowers, and cauline leaves because of their low counts compared to samples from the roots and larger phyllosphere tissues. Stem and silique samples from the flowering stage were also excluded because of low sample counts compared to samples from those tissues during senescence.

To construct networks, samples were grouped into three plant developmental stages: vegetative, flowering, and senescent. Samples in the study were collected from 2-leaf rosettes, 4-leaf rosettes, 6-leaf rosettes, 8-leaf rosettes, flowering plants, and senescent plants. Since the earlier vegetative stages yielded a relatively small number of samples, only the latest rosette stage sampled (6-leaf in Year 2 and 8-leaf in Year 1) was used to assemble the “vegetative” networks.

ASV prevalence and abundance patterns varied between plant tissues and developmental stages as well as across field sites and between study years (2). Root, rosette leaf, stem, and silique samples were therefore considered independently. For each tissue type, samples from different developmental stages and sites were used to infer independent networks of interactions among only the ASVs counted during both study years. Before network inference, the count tables were pruned to remove ASVs present in fewer than three samples and with fewer than two hundred and fifty counts. These quantities were settled upon by testing different thresholds for ASV prevalence and count totals in the dataset. After network inference, node degree and betweenness centrality were plotted against prevalence and count total. A large number of very highly connected nodes with low prevalence or abundance indicated the likely presence of spurious edges in the network connected to the rare ASVs. Thus, the edges in the networks are based on at least three observations totaling at least two hundred and fifty counts across two years of data collection.

Network inference

Networks of ASV relationships were inferred with both an inverse covariance and a correlation-based approach.

Interactions were first inferred with the SPIEC-EASI (**SP**arse **I**nvers**E** **C**ovariance Estimation for **E**cological **A**ssociation **I**nference) pipeline (9). Conditional dependencies between ASVs were found with an inverse covariance matrix produced by log-ratio transformation of counts. Then, the inferred relationships between ASVs were represented in a graph constructed by neighborhood selection. The appropriate sparseness for the graph was found with the Stability Approach to Regularization Selection (StARS) method, based on random subsampling of the graph (10). Weights were assigned to the edges of the resulting undirected graph based on coefficients for the Pearson correlation between node abundances. [SPIEC-EASI parameters list: method="mb", lambda.min.ratio=1e-1, rep.num=100]

Relationships among ASVs were next inferred with the SparCC algorithm (11) within the R SPIEC-EASI package. Two-sided empirical p-values were calculated for the correlations inferred with SparCC. Correlations were filtered for significance at α = 0.001 and coefficient absolute value > 0.3. The matrix of filtered correlation coefficients was then used to produce an undirected graph with edges weighted by the absolute value of the coefficients.

[SparCC parameters list: iter=20,inner_iter=10, R=10, pval.sparccboot(sided="both")]

Network attributes

The ASV count table for each set of conditions (site, stage, and tissue) was pruned to the ASVs present in the interaction network for those conditions. For each ASV, the following properties were calculated from the count tables and assigned to the corresponding node in the network: mean raw abundance (average counts per sample), mean relative abundance (average fractional abundance), and prevalence (number of samples in which present). Each ASV was also assigned taxonomy at the levels of phylum, class, order, and family.

The phylogenetic distances spanned by network edges were determined as follows: First, all possible pairwise combinations of ASVs in the 16S gene tree for a set of conditions (site, stage, and tissue) were found. Second, the total branch length on the tree between each ASV combination was calculated and stored in a dataframe with the names of the ASVs. Third, the dataframe with branch lengths was used as an edge list to construct an undirected graph with phylogenetic distance as an edge attribute. The intersection of this graph and the inferred interaction network produced a graph with phylogenetic distance as an edge attribute. A linear model was fit to the absolute values of edge correlation coefficients vs. phylogenetic distances to assess whether the ecological and evolutionary relationships between bacteria might be related.

Analysis of network properties

Before analysis, unattached nodes (with degree zero) were removed from networks. The number of connected nodes (bacterial ASVs), number of edges (inferred ASV interactions), and the fraction of edges with positive correlations between the ASV abundances (positive interactions) were tallied for each network.

The extent of structure in each interaction network was assessed by clustering the graph into communities based on edge betweenness (igraph's cluster_edge_betweenness) and calculating modularity with the resulting community structure (igraph's modularity). Since modularity depends on network size, this property was compared between networks with z-scores of modularity in distributions based on one hundred randomly generated graphs (igraph's erdos.renyi.game) with the same number of nodes and edges as the empirical network.

The degree distribution of each network was fit to a power law to assess whether it was consistent with that expected for a scale-free network (12). A null degree distribution was generated for each network with the Barabási-Albert model for growth with preferential attachment using the “psumtree” algorithm in igraph's sample_pa function and a number of nodes equivalent to that in the empirical network.

A local clustering coefficient was calculated for each node using igraph's transitivity function. A linear model was fit to the log10-transformed clustering coefficient vs. log10 degree for each node to assess whether the network showed evidence of a hierarchical structure (13).

All network properties were compared between networks from different tissues, sites, and stages as well as between networks generated from the same ASV counts with different methods.

Identifying hubs

Given that global network properties changed in datasets spanning plant development, hubs were detected with a dynamic threshold rather than a single cutoff for node degree. Both node degree (number of edges attached to a node) and betweenness centrality (the number of shortest paths going through a node) were used to find hubs, following the approach by Agler et al 2016 (14). Nodes with degree and betweenness z-scores corresponding to the ninetieth percentile of a network were classified as hubs. The conservation of edges and hubs across networks were compared using heatmaps and the UpSetR package (15).

(2) Beilsmith K, Perisin M, Bergelson J. Natural bacterial assemblages in *Arabidopsis thaliana* tissues become more distinguishable and diverse during host development.

(3) R Core Team. R: A language and environment for statistical computing. 2018. R Foundation for Statistical Computing, Vienna, Austria. <https://www.R-project.org/>.

(4) McMurdie PJ, Holmes S. phyloseq: an R package for reproducible interactive analysis and graphics of microbiome census data. PloS one. 2013;8(4). doi: [10.1371/journal.pone.0061217](https://doi.org/10.1371/journal.pone.0061217)

(5) Csardi G, Nepusz T. The igraph software package for complex network research. InterJournal, complex systems. 2006 Jan 11;1695(5):1-9. <https://igraph.org/r>

(6) Jari Oksanen, F. Guillaume Blanchet, Michael Friendly, Roeland Kindt, Pierre Legendre, Dan McGlinn, Peter R. Minchin, R. B. O'Hara, Gavin L. Simpson, Peter Solymos, M. Henry H. Stevens, Eduard Szoecs and Helene Wagner (2017). vegan: Community Ecology Package. R package version 2.4-5. <https://CRAN.R-project.org/package=vegan>

(7) H. Wickham. ggplot2: Elegant Graphics for Data Analysis. Springer-Verlag New York, 2016.
