## Supplementary Figures for "Transient interactions and influence among bacteria in field-grown *Arabidopsis thaliana* tissues"

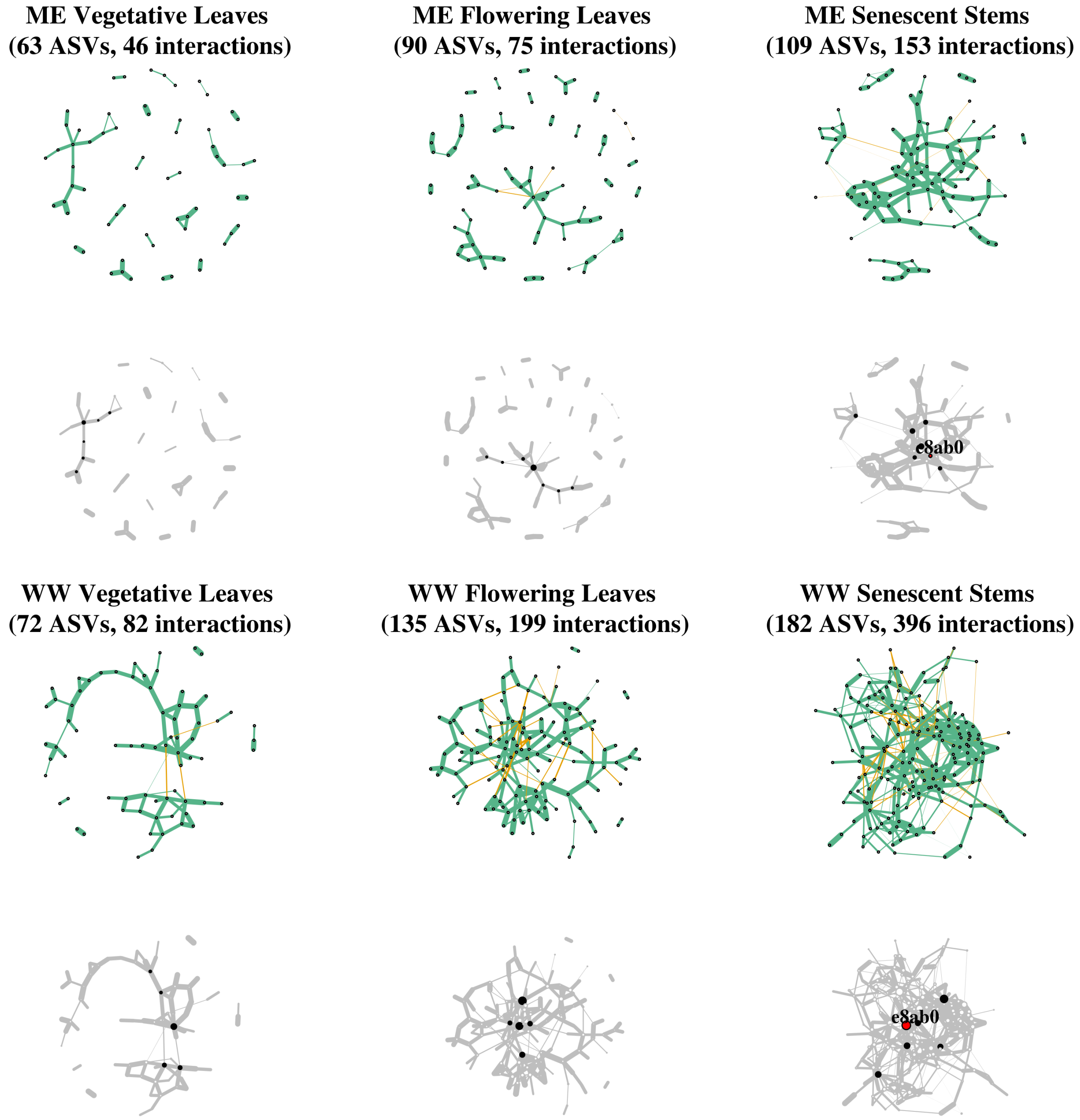


<u>Figure S1</u>

These phyllosphere bacterial networks further support points made with the root networks in Figure 1A. Color indicates positive (green) and negative (orange) abundance correlations and line thickness corresponds to the correlation coefficient. Hubs are highlighted in black and those conserved across networks are red and labeled with partial ASV identifiers. Only one ASV (e8ab0), in the genus *Nocardioides*, was conserved.

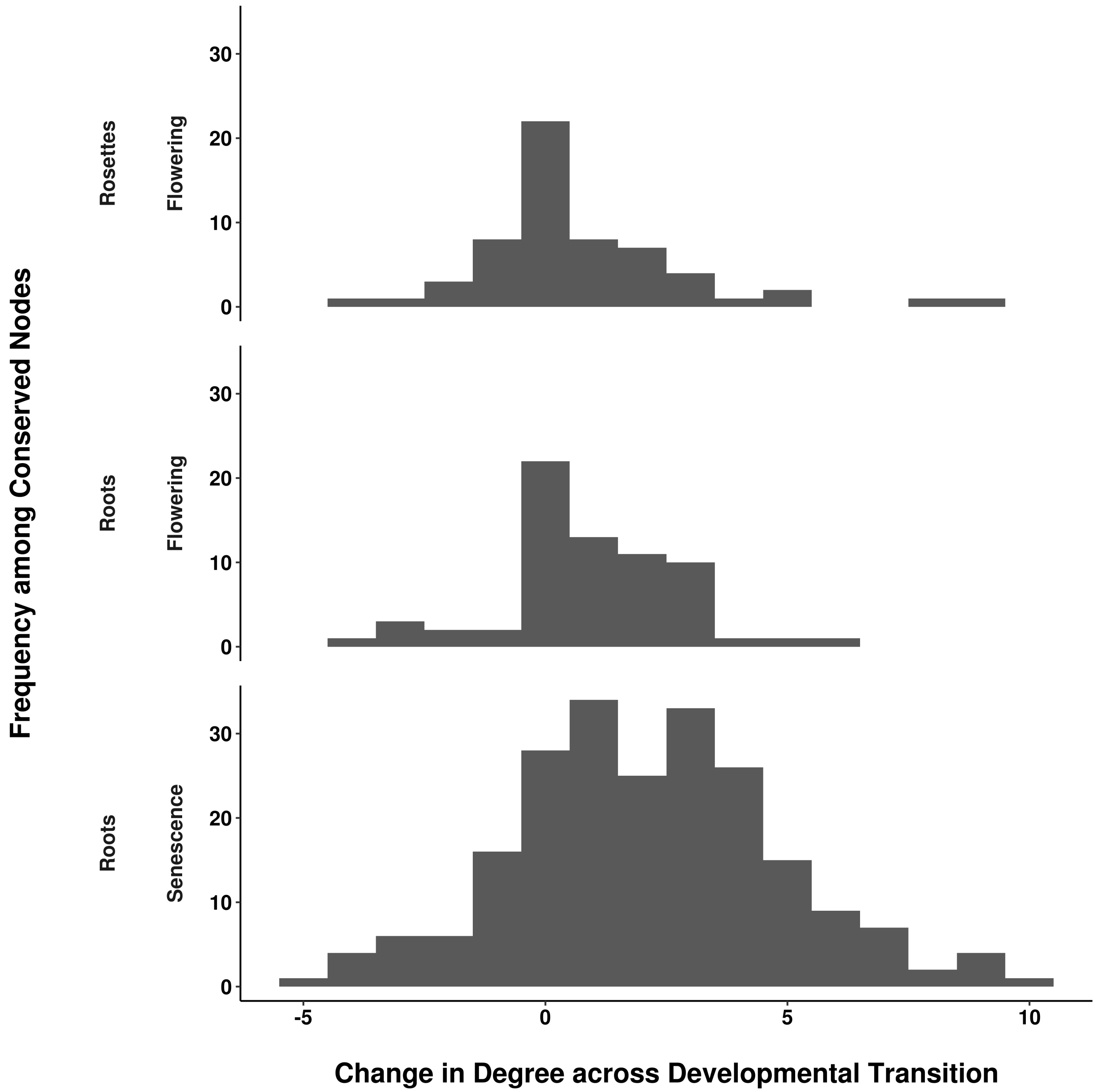


Figure S2

The number of ASV interaction partners tends to increase over major developmental transitions. For ASVs (nodes) remaining after a transition, the number of edges (degree) before and after the transition were compared. On the x-axis, positive numbers indicate net connections gained while negative numbers indicate net connections lost. The bar height on the y-axis indicates the frequency of conserved nodes with the corresponding number of gains or losses. Interactions gained and lost are shown for the transition to flowering in roots and rosette leaves and for senescence in roots (vertical panels).

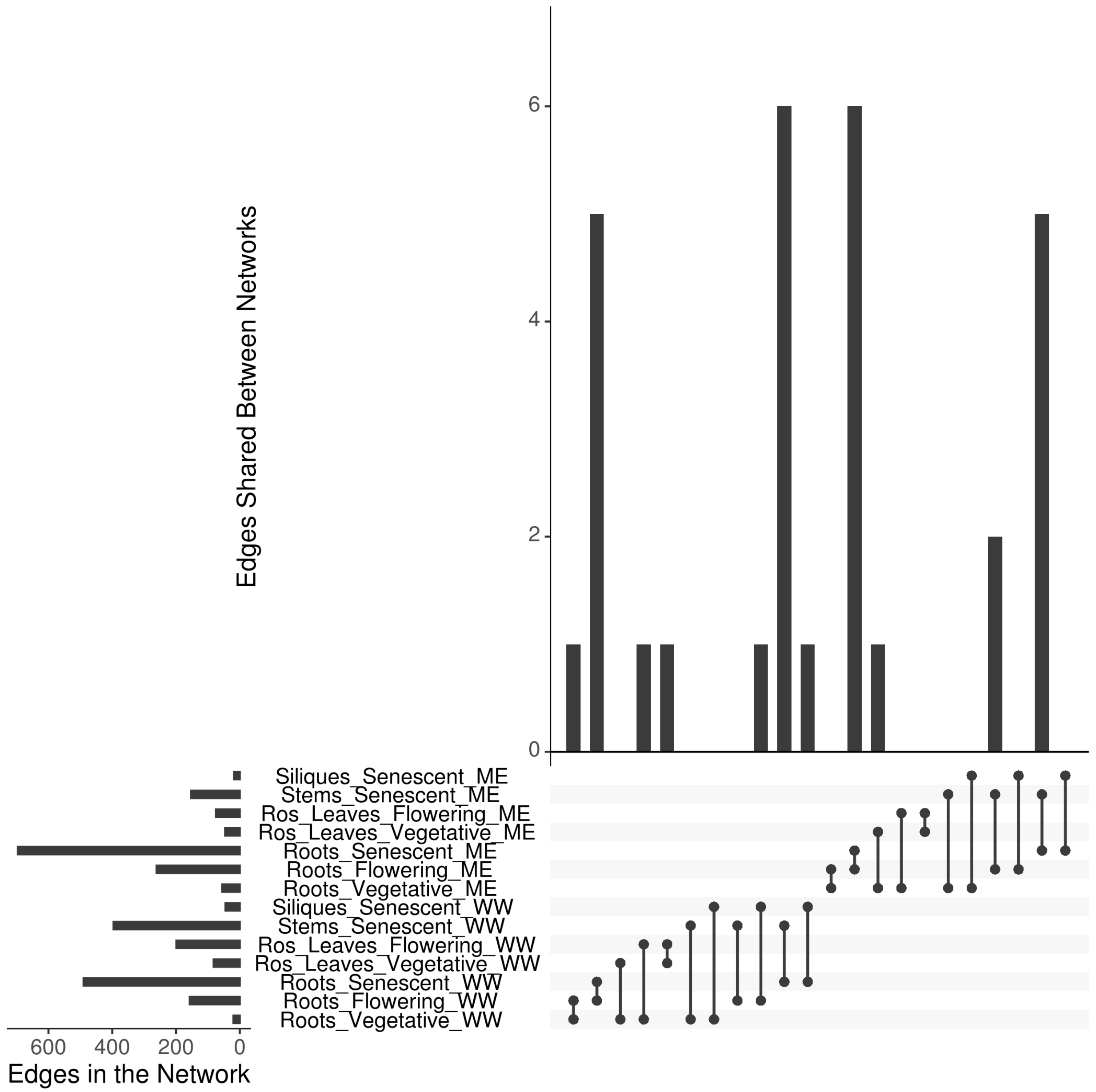


Figure S3

Interactions in the networks are transient, as evidenced by the low number of edges conserved between ASVs that occur before and after major developmental transitions. Network comparisons are indicated on the x-axis by filled circles connected by lines. The number of overlapping edges in each comparison is shown on the y-axis. The total number of edges in each network is plotted to the left of the network name.

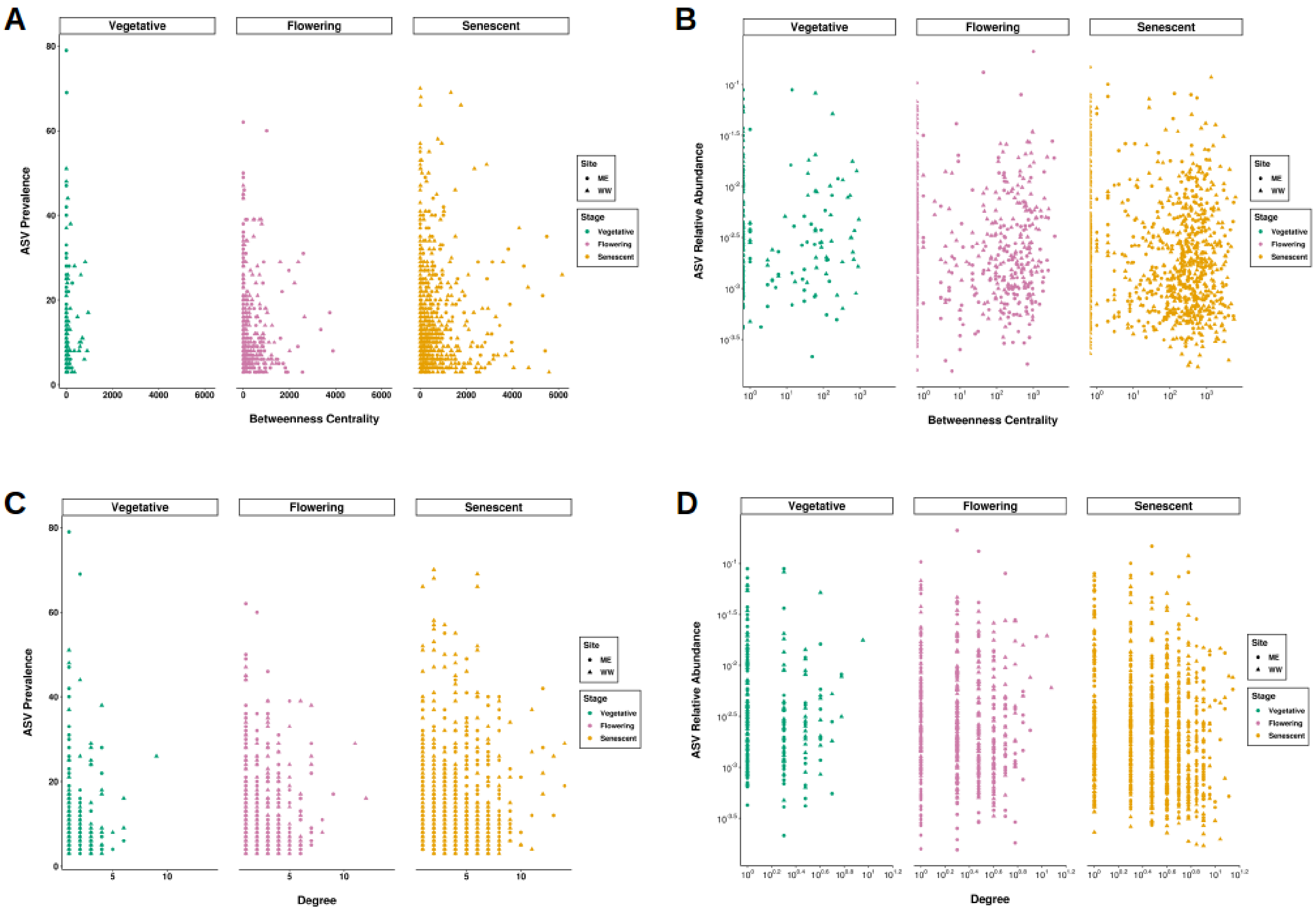


<u>Figure S4</u>

Hubs were identified by degree and betweenness centrality. These criteria for influence did not correlate with ASV prevalence (the number of samples in which the ASV was present) or raw abundance (the total number of counts for the ASV).

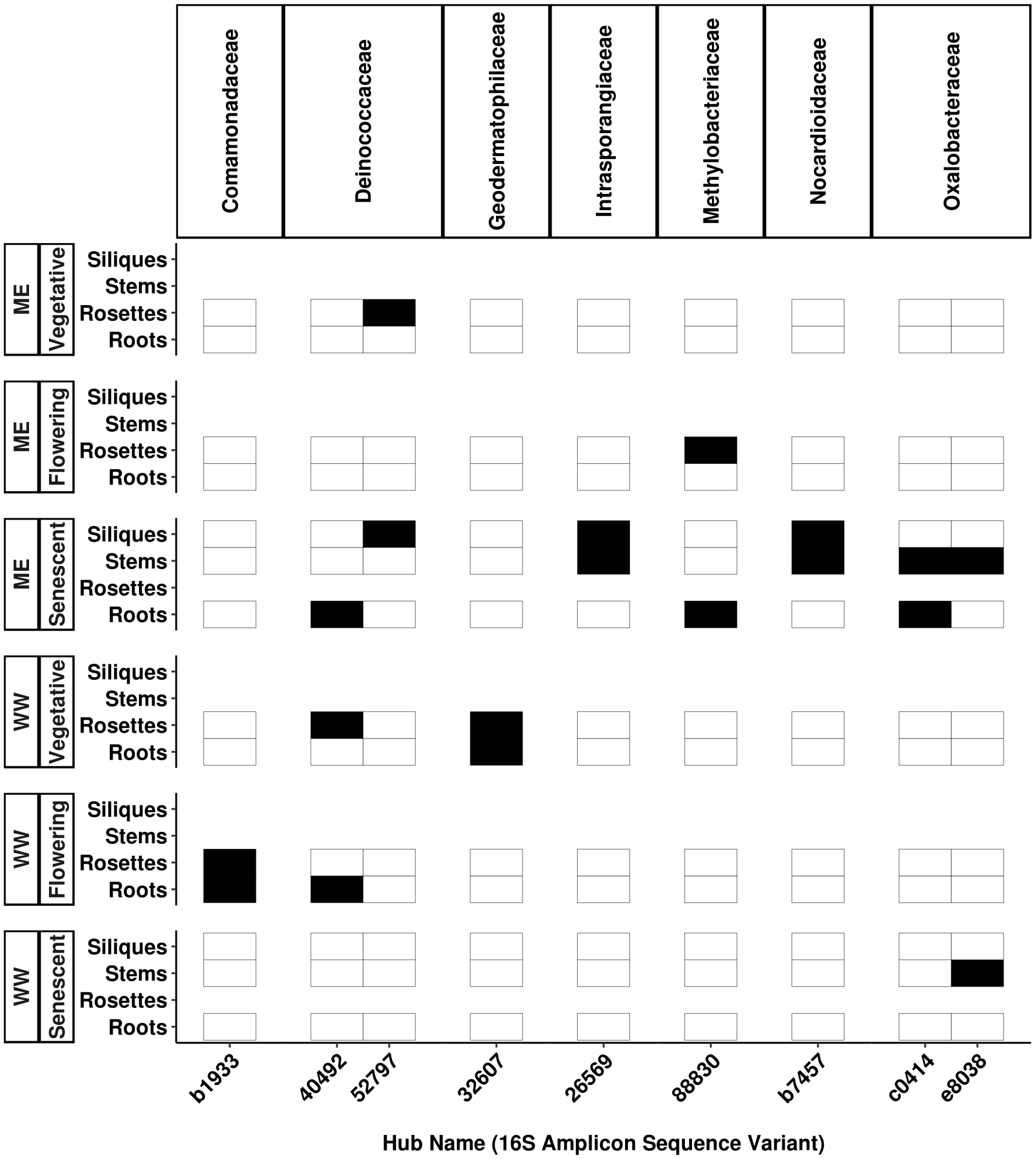


Figure S5

Distributions of recurring hubs (columns) in inverse covariance networks. Hubs are organized by family and compared across developmental stages, field sites (vertical panels) and tissues (rows). Black cells indicate the ASV is both present and meets the hub criteria; white cells indicate the ASV is either not present or not a hub.

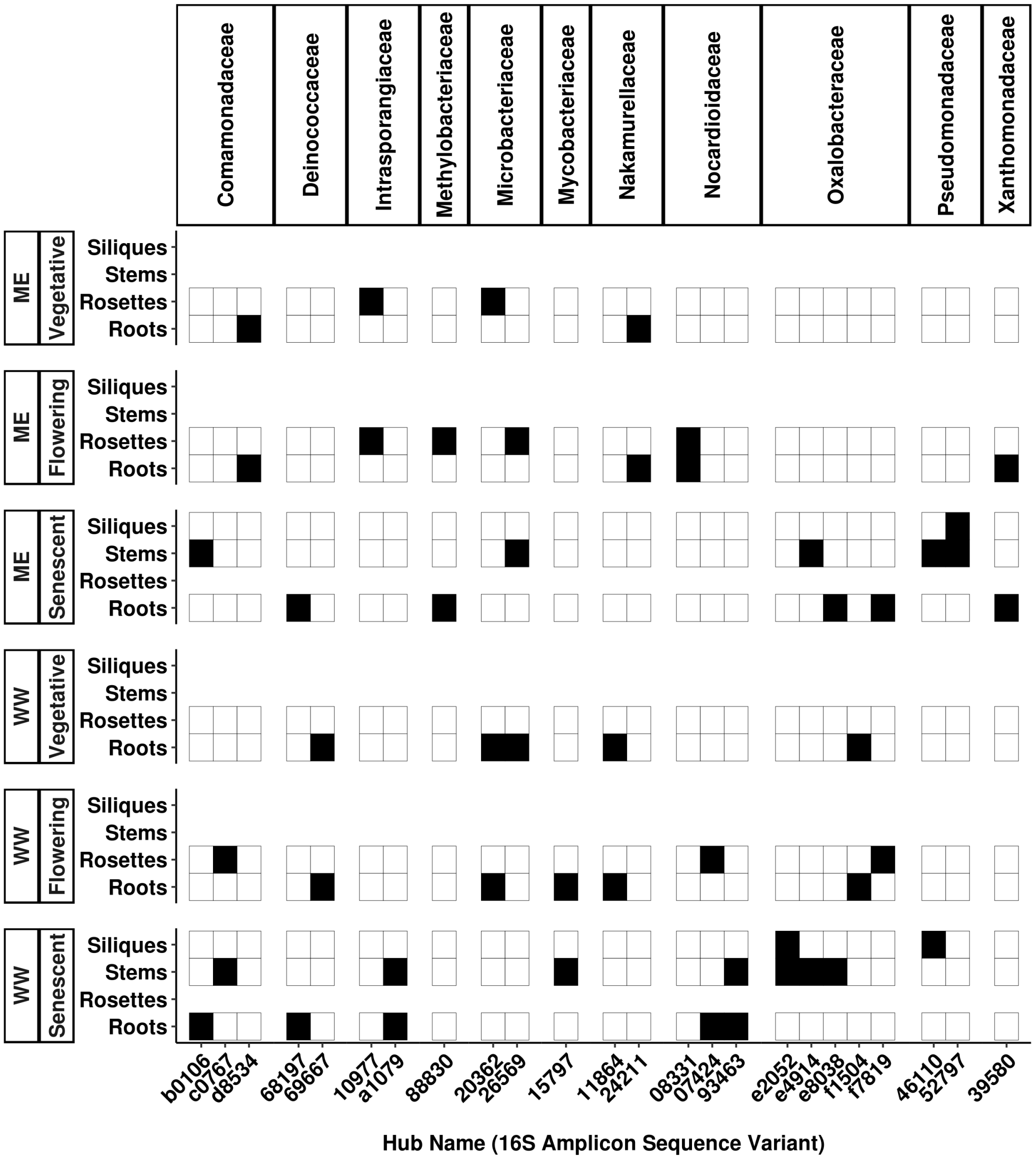


<u>Figure S6</u>

Distributions of recurring hubs (columns) in correlation-based networks. Hubs are organized by family and compared across developmental stages, field sites (vertical panels) and tissues (rows). Black cells indicate the ASV is both present and meets the hub criteria; white cells indicate the ASV is either not present or not a hub.
